# A toolkit for converting Gal4 into LexA and Flippase transgenes in *Drosophila*

**DOI:** 10.1101/2022.12.27.522021

**Authors:** Sasidhar Karuparti, Ann T. Yeung, Bei Wang, Pedro F. Guicardi, Chun Han

**Author notes:** These authors contributed equally to this work (Sasidhar Karuparti, Ann T. Yeung).

## Abstract

*Drosophila* has been a powerful model system for biological studies due to the wide range of genetic tools established for it. Among these tools, Gal4 is the most abundant, offering unparalleled tissue- and developmental stage-specificity for gene manipulation. In comparison, other genetic reagents are far fewer in choices. Here we present a genetic toolkit for converting Gal4 strains into LexA and Flippase transgenes through simple genetic crosses and fluorescence screening. We demonstrate the proof-of-principle by converting ten Gal4 lines that exhibit diverse tissue specificities and examined the activity patterns of the converted LexA and Flippase lines. Gal4-to-LexA and Flp conversion is fast and convenient and should greatly expand the choices of LexA and Flp for binary expression and FRT-based mosaic analysis, respectively, in *Drosophila*.

## Introduction

*Drosophila* is a powerful model system for studying developmental biology, cell biology, neurobiology, and genetics. This power largely lies in the numerous genetic tools available in *Drosophila* for manipulating the genome and gene activity. Commonly used tools include binary gene expression systems (Brand and Perrimon 1993; Lai and Lee 2006; Potter *et al*. 2010), site-specific recombinases (Golic and Lindquist 1989; Bischof *et al*. 2007; Nern *et al*. 2011), clustered regularly-interspaced palindromic repeats (CRISPR)/Cas9 (Gratz *et al*. 2013; Ewen-Campen *et al*. 2017; Bosch *et al*. 2020; Bosch *et al*. 2021), and many more. Binary expression systems allow for expression of transgenes in spatially and temporally restricted, and developmental stage-specific manners. Their variants (Osterwalder *et al*. 2001; Luan *et al*.2006) and modifiers (McGuire *et al*. 2003) further increase the precision of the control and offer greater flexibility. Site-specific recombination systems enable rearrangement of genomic DNA and allow for development of sophisticated methods for creating genetic mosaics (Dang and Perrimon 1992; Struhl and Basler 1993; Xu and Rubin 1993; Lee and Luo 1999). More recently, CRISPR/Cas9 tools allow convenient generation of permanent or tissue-specific mutations (Gratz *et al*. 2013; Port *et al*. 2014; Poe *et al*. 2017), replacement of endogenous genomic sequences with desired ones (Gratz *et al*. 2014; Port *et al*. 2014), and insertion of exogenous sequences at precise loci (Lee *et al*. 2018).

Since the introduction of the yeast Gal4/UAS system into *Drosophila*(Brand and Perrimon 1993), tens of thousands of Gal4 strains have been generated using diverse approaches (Brand and Perrimon 1993; Sharma *et al*. 2002; Pfeiffer *et al*. 2008; Jenett *et al*. 2012; Kvon *et al*. 2014). In each strain, the transcription factor Gal4 is expressed under the control of specific enhancer elements and thus exhibits a unique expression pattern. This vast Gal4 resource makes investigations of gene function feasible in virtually any tissue and at any developmental stage. In comparison, the availability of other genetic tools is much more limited, hampering researchers’ ability to use orthogonal approaches. For example, LexA/LexAop is another popular binary system (Lai and Lee 2006), but the limited choices for LexA make the system less flexible to use as compared to Gal4. Flp/FRT is a site-specific recombination system; it enables mosaic analysis techniques such as flp-out (Struhl and Basler 1993) and MARCM (Lee and Luo 1999). Although temporally inducible Flp is available for these techniques, tissuespecific Flp would greatly simplify and streamline large-scale applications such as genetic screens (Huang *et al*. 2014; Neukomm *et al*. 2014). However, tissue-specific Flp resources are also very limited at present. Thus, convenient methods for generating additional tissue-specific LexA and Flp lines will be greatly beneficial to the *Drosophila* research community.

CRISPR/Cas9 provides an attractive option for converting existing Gal4 resources into other systems. It has been widely used in *Drosophila* for precise genome engineering (Gratz *et al*. 2014; Port *et al*. 2014), which takes advantage of double-strand break (DSB)-induced DNA repair through the homologous recombination pathway. While it is common to perform gene replacement through embryo injections, the recently developed homology assisted CRISPR knock-in (HACK) method demonstrated the feasibility of converting existing Gal4 lines into other tissue-specific reagents through simple genetic crosses (Lin and Potter 2016). This method eliminates the need for injection because the necessary genetic components are brought together by genetic crosses to induce homologous recombination in the fly germline. This method has already been successfully used to convert Gal4 strains into tissue-specific QF, split-Gal4, Gal80, and Cas9 lines (Lin and Potter 2016; Xie *et al*. 2018; Chen *et al*. 2020; Koreman *et al*. 2021). Although HACK has the potential to greatly expand available genetic resources for researchers, this method has not been used to make LexA or Flp reagents, which would be useful complementary tools to the ones previously made.

In this study, we developed tools that allow conversion of Gal4 lines into LexA and Flp lines based on HACK. We demonstrate the proof-of-principle by converting several Gal4 drivers that are expressed in stem cells, epithelial cells, muscles, adipocytes, glia, and neurons. We show that the tissue-specificity of these LexA and Flp reagents is maintained. This method is convenient and can be applied at a large scale for rapid expansion of LexA and Flp resources.

## Materials and Methods

### Fly strains

The fly strains used in this study are listed in the Reagent Table.

### Construction of HACK donor vectors

The HACK donor vectors were constructed by modifying pHACK(Gal4)-DONR(T2A-Cas9) (Addgene # 194768), a donor vector for converting Gal4 into Cas9 (Koreman *et al*. 2021). The homology arms (HAs) are 1119 bp for 5’ and 1194 bp for 3’. We use two gRNAs targeting the middle of Gal4 in the donor vector to increase the CRISPR efficiency. To make pHACK(Gal4)-DONR(T2A-LexAGAD), a nlsLexAGAD partial sequence was PCR amplified from pDEST-APIC-LexAGAD (Poe *et al*. 2017) using oligos GAAGCGGAGGCgctagcATGCCACCCAAGAAGAAGC and CACATATAGGACTTTTTTCTGCAGTCACTGTCTTATCCAGCTC. The fragment was assembled with Nhel/PstI digested pHACK(Gal4)-DONR(T2A-Cas9) through NEBuilder HiFi DNA Assembly. To make pHACK(Gal4)-DONR(T2A-Flp1), the Flp1 CDS was PCR amplified from pDEST-APIC-Flp1 (Poe *et al*. 2017) and assemble with Nhel/Agel digested pHACK(Gal4)-DONR(T2A-Cas9) through NEBuilder HiFi DNA Assembly.

### 13XLexAop2-GFPnls-PEST

A DNA fragment containing SV40 nuclear localization signal (nls) and a protein destabilization PEST signal from mouse ornithine decarboxylase (NP_038642.2; corresponding to aa 423-461) was synthesized (Integrated DNA Technologies, Inc) and cloned into pAPLO (Poe *et al*. 2017). The superfolder GFP (sfGFP) coding sequence was PCR-amplified from pIHEU-AV-sfGFP (Sapar *et al*.2018), with syn21 (a translation enhancer), start codon, and SV40 nuclear localization signal (nls) in the forward primer, and cloned in-frame before SV40nls and PEST in pAPLO.

### Generation of transgenes

Injections were carried out by Rainbow Transgenic Flies (Camarillo, CA 93012 USA) to transform flies through φC31 integrase-mediated integration into attP sites. Each HACK donor vector was inserted into the attP40 (on the 2^nd^ chromosome) and attP^VK00027^ (on the 3^rd^ chromosome) sites. 13XLexAop2-GFPnlsPEST was inserted into attP^VK00033^ site.

### Conversion of Gal4 to LexAGAD and Flp

Conversion experiments were conducted similarly to Figure 2. A germline-specific nos-Cas9 (Port *et al*. 2014) or *Bam-Cas9* (Chen *et al*. 2020) on the X chromosome was combined with the appropriate donor transgene and a Gal4 insertion into the same fly through two sequential crosses. The Gal4 and the donor transgene were located on two homologous chromosomes. Founder flies containing all three components were crossed to reporter lines. For *nos-Cas9*, female founders appeared to have higher efficiencies of conversion than male founders. For *Bam-Cas9*, we used male founders because *Bam-Cas9* was reported to have higher activity in the male germline (Chen *et al*. 2020). For Flp conversion, *10XUAS-IVS-mCD8::RFP 13XLexAop2-mCD8::GFP; CoinFLP-LexA::GAD.GAL4* (BDSC # 58754) was initially used as the reporter. Later, *Tub>STOP>LexAGAD::VP16; 13XLexAop2-6XGFP* was built as a more convenient reporter. For LexAGAD conversion, *13XLexAop2-6XGFP* (Shearin *et al*. 2014) was used as the reporter. 3^rd^ instar larvae showing the expected GFP expression patterns were screened from the progeny under a Nikon SMZ18 fluorescence stereomicroscope and recovered for development into adulthood. The resulting flies were crossed to proper balancer stocks to separate the reporter chromosome and the converted LexAGAD or Flp chromosome. In our hands, it takes approximately 60 days from the beginning to the establishment of a converted line. A subset of the converted LexAGAD and Flp lines were validated by genomic PCR (Figure S1).

**Figure 1.**
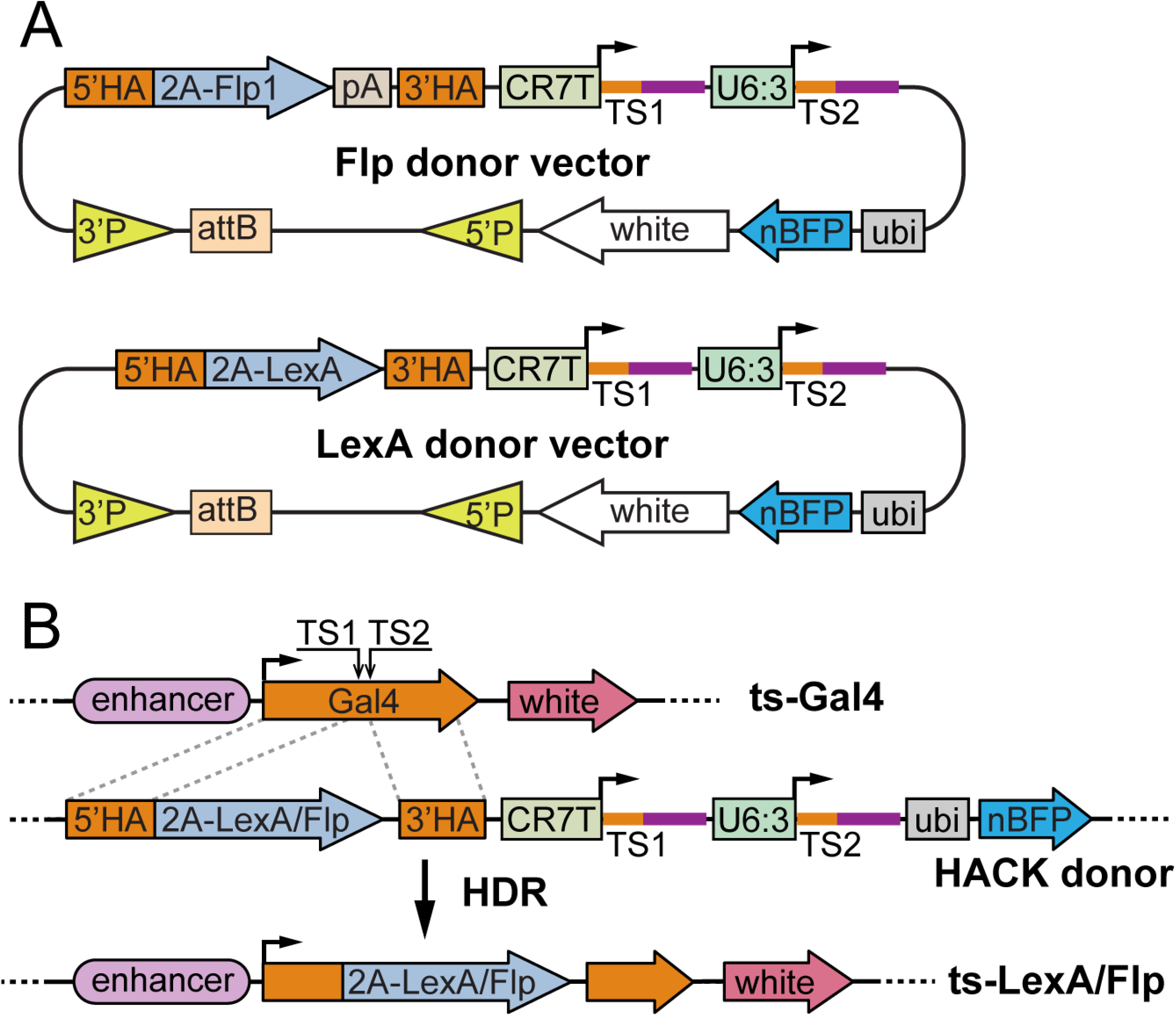
Construct designs and the principle of conversion. (A) Diagrams of Flp and LexA HACK donor constructs. The vectors were constructed in pAC, a dual-transformation backbone (via PhiC31 and P-transposase) that carries a mini-white selection marker. See text for descriptions of other components. pA: polyA tail; TS: gRNA target sequence; P: P-element. (B) Diagram of Gal4-to-Flp/LexA conversion using a HACK donor line. The donor expresses two gRNAs (TS1 and TS2) targeting the tissue-specific (ts) Gal4, which results in in-frame incorporation of 2A-Flp/LexA into the Gal4 locus through homology-directed repair (HDR). The donor expresses *ubi-nBFP* that can be selected against when screening for convertants.

**Figure 2.**
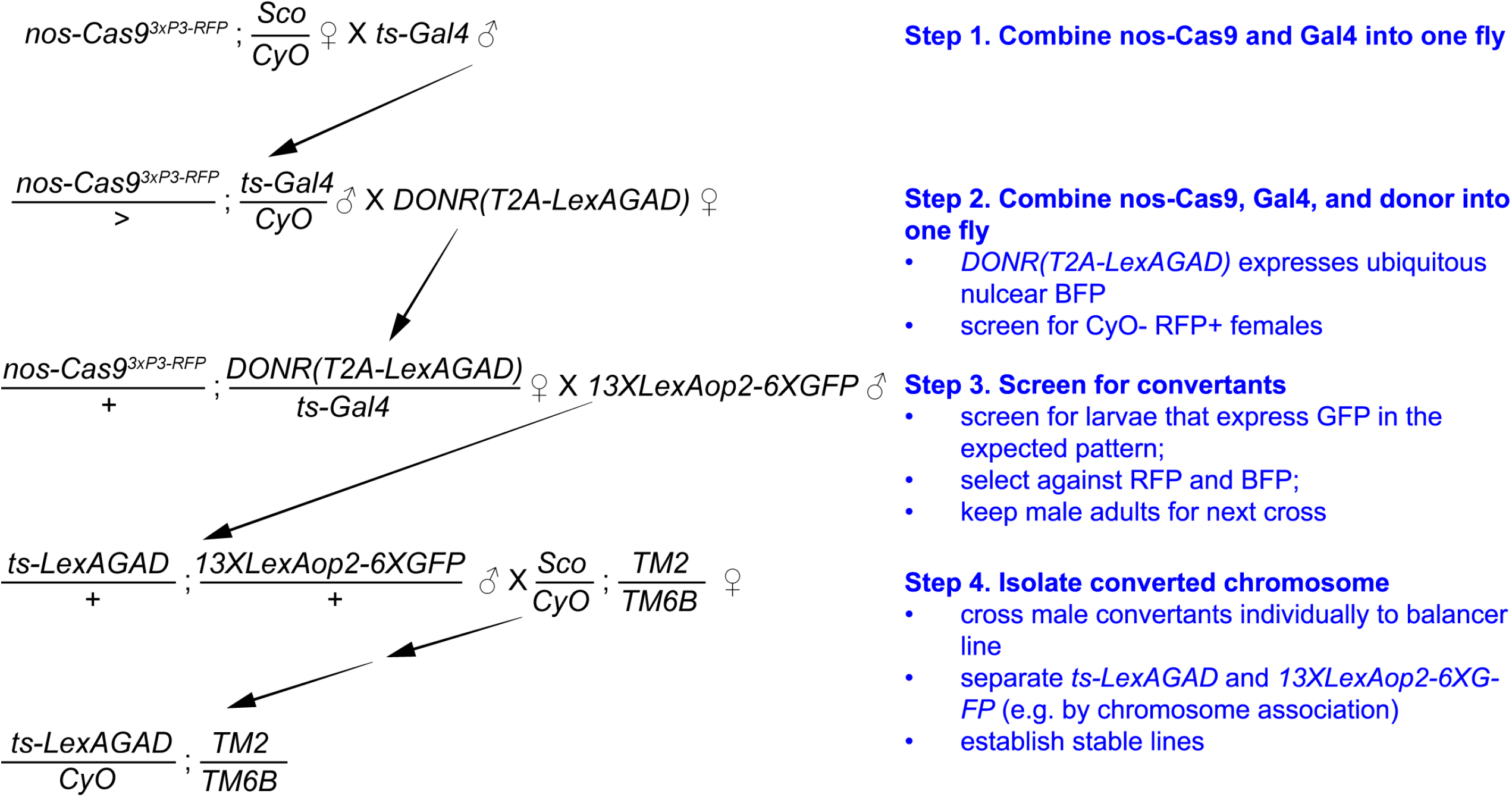
Example crossing scheme for converting Gal4 into LexAGAD. Illustrated is a crossing scheme for converting a second-chromosome Gal4 line into LexAGAD line. The most critical step is the screening of convertants based on fluorescence in the expected pattern (Step 3). 50-300 larvae, depending on the Gal4, usually need to be screened to get enough convertant candidates (5-10 larvae). This particular example utilizes *nos-Cas9* as the germline Cas9 and a donor transgene on the second chromosome. *nos-Cas9* is more effective in female founders than in males. *nos-Cas9* can be substituted by *bam-Cas9-P2A-Flp*, which is more effective in male founders than in females. The donor transgene can also be on a nonhomologous chromosome.

### Validation of expression pattern/imaging

The converted LexAGAD and Flp lines were crossed to GFP reporter lines according to Table 1. GFP expression patterns in wandering 3^rd^ instar larvae were examined with a Leica SP8 confocal equipped with a 20X oil objective. For brain expressions, we dissected larval brains and stained the samples with the primary antibody NC82 (Developmental Studies Hybridoma Bank, 1:100) and the secondary antibody Cy5 donkey anti-mouse antibody (Jackson ImmunoResearch Laboratories; 1:400) to visualize neuropiles. For wing disc expression, we dissected larvae and stained the samples with 4’,6-diamidino-2-phenylindole (DAPI; 1:36000). For all other crosses, we imaged the body walls of live larvae.

**Table 1.**
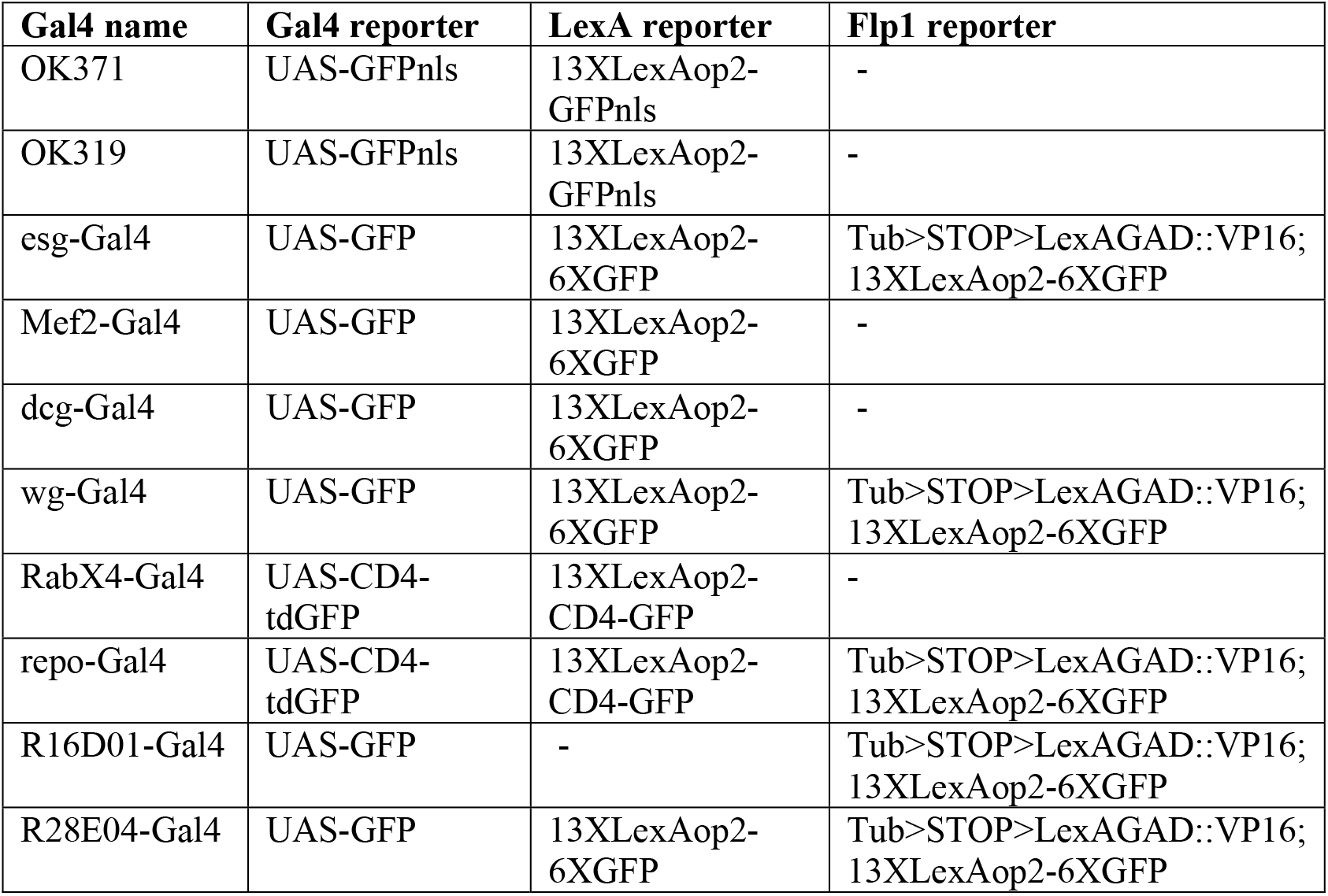
Crosses for validating Gal4, LexAGAD, and Flp activity patterns.

## Results

### Construct designs and the principle of conversion

To enable conversion of Gal4 lines into LexA and Flp lines, we generated two HACK donor transgenic constructs (Figure 1A), building on a dual-gRNA vector we previously optimized for CRISPR-mutagenesis in the *Drosophila* germline (Koreman *et al*. 2021). Each donor construct carries three functional units that collectively enable homology-directed repair (HDR)-mediated conversion and larval screening. First, a gRNA cassette encodes two gRNAs driven by two separate polymerase III promoters (CR7T and U6:3) to target the Gal4 coding sequence. The gRNAs adopt the gRNA2.1 scaffold (Grevet *et al*. 2018), which is more efficient than the original and the gRNA(F+E) scaffolds in mutagenesis in mammalian cells and *Drosophila*(Grevet *et al*. 2018; Koreman *et al*. 2021). We used two, instead of only one, gRNAs to increase the possibility of DNA double-strand break. Second, a donor sequence contains the coding sequence of 2A-Flp or 2A-LexA flanked by two homology arms (HAs) from the Gal4 coding sequence. While the DNA binding domain of LexA is fused in-frame with the Gal4 activation domain (GAD, within the 3’ HA) in the LexA donor construct, the Flp sequence is followed by a *hsp70* polyA for transcription termination in the Flp donor construct. Third, a nuclear BFP (nBFP) marker driven by a polyubiquitin promotor (ubi) serves as a selection marker for distinguishing the donor chromosome. The donor vectors were constructed in pAC (attB-CaSpeR), a backbone that is compatible with both P-element- and PhiC31-mediated transformation (Han *et al*. 2011).

The conversion of Gal4 is induced in the *Drosophila* germline by combining the LexA or Flp donor transgene, a Gal4 of interest, and a germline specific Cas9 (such as *nos-Cas9 and bam-Cas9-P2A-Flp*) (Figure 1B). gRNA/Cas9 produces DSBs in the middle of the Gal4 coding sequence. Homology-directed repair of the DSBs using the donor sequence as a template will result in in-frame incorporation of 2A-LexA or 2A-Flp in the original Gal4 locus. During translation, the “self-cleaving” 2A peptide releases a truncated and nonfunctional Gal4 and LexAGAD or Flp as two separate proteins. Thus, the expression pattern of the resulting LexAGAD/Flp line should reflect that of the original Gal4.

The conversion can be carried out through several simple steps of genetic crosses (illustrated in Figure 2 for converting Gal4 insertions on the 2^nd^ chromosome to LexA versions). Successful conversion events will result in chromosomes that carry tissue-specific LexA or Flp and can be identified using specific LexA or Flp reporters. We used *13XLexAop2-6XGFP* (Shearin *et al*. 2014) as a LexA reporter and *10XUAS-IVS-mCD8::RFP 13XLexAop2-mCD8::GFP; CoinFLP-LexA::GAD.GAL4* and *Tub>STOP>LexAGAD::VP16; 13XLexAop2-6XGFP* as Flp reporters. These reporters express high levels of fluorescent proteins in the Flp/LexA expressing tissues, making it easy to identify larvae carrying the converted chromosome, even when the expression domain is restricted. Although our method was designed for converting Gal4 lines that exhibit recognizable expression patterns in the whole larva, similar approaches should allow conversion of Gal4 lines that show visible expression patterns in adults.

### Conversion of example Gal4 lines

We inserted each donor construct into two attB sites, one on the second chromosome and the other on the third, through PhiC31-mediated integration. To test the effectiveness of the conversion, we chose 10 Gal4 lines that show tissue-specific expression in the larva (Table 2). These Gal4 transgenes are at various locations on the second and the third chromosomes and were created by diverse means, including enhancer trap (Brand and Perrimon 1993), enhancer fusion with random insertion (Ranganayakulu *et al*. 1998), enhancer fusion with targeted insertion (Pfeiffer *et al*. 2008), and recombineering of genomic DNA clones followed by targeted insertion (Chan *et al*. 2011). These Gal4s are controlled by regulatory sequences from different genes and are expressed in diverse larval cell types, including epithelial cells, muscles, neurons, glia, adipocytes, hemocytes, and stem cells.

**Table 2.**
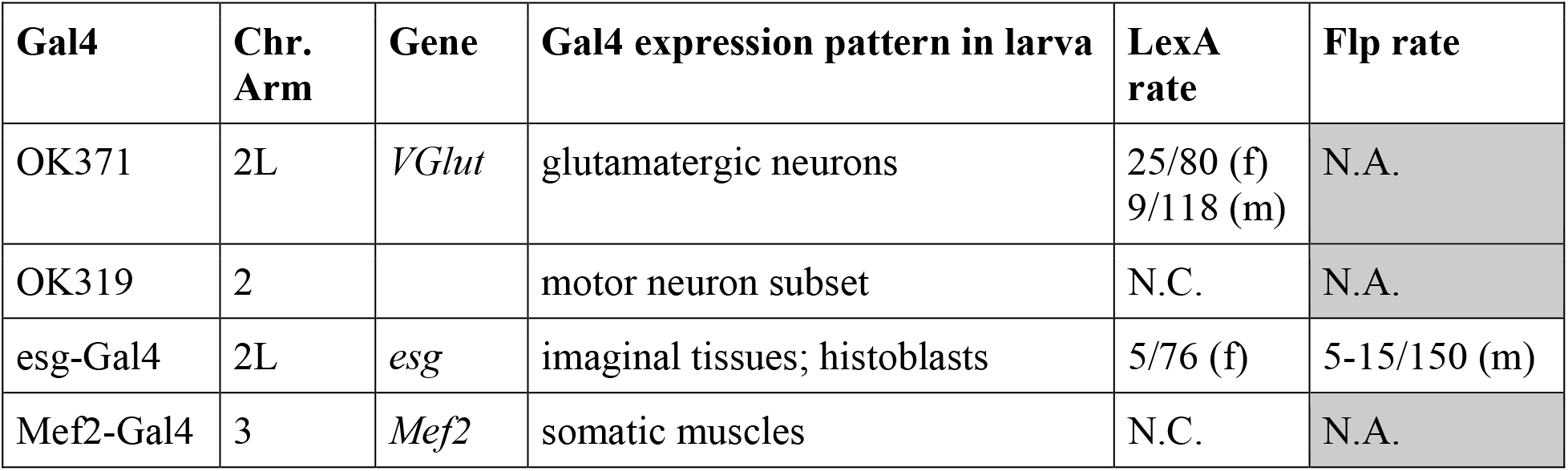
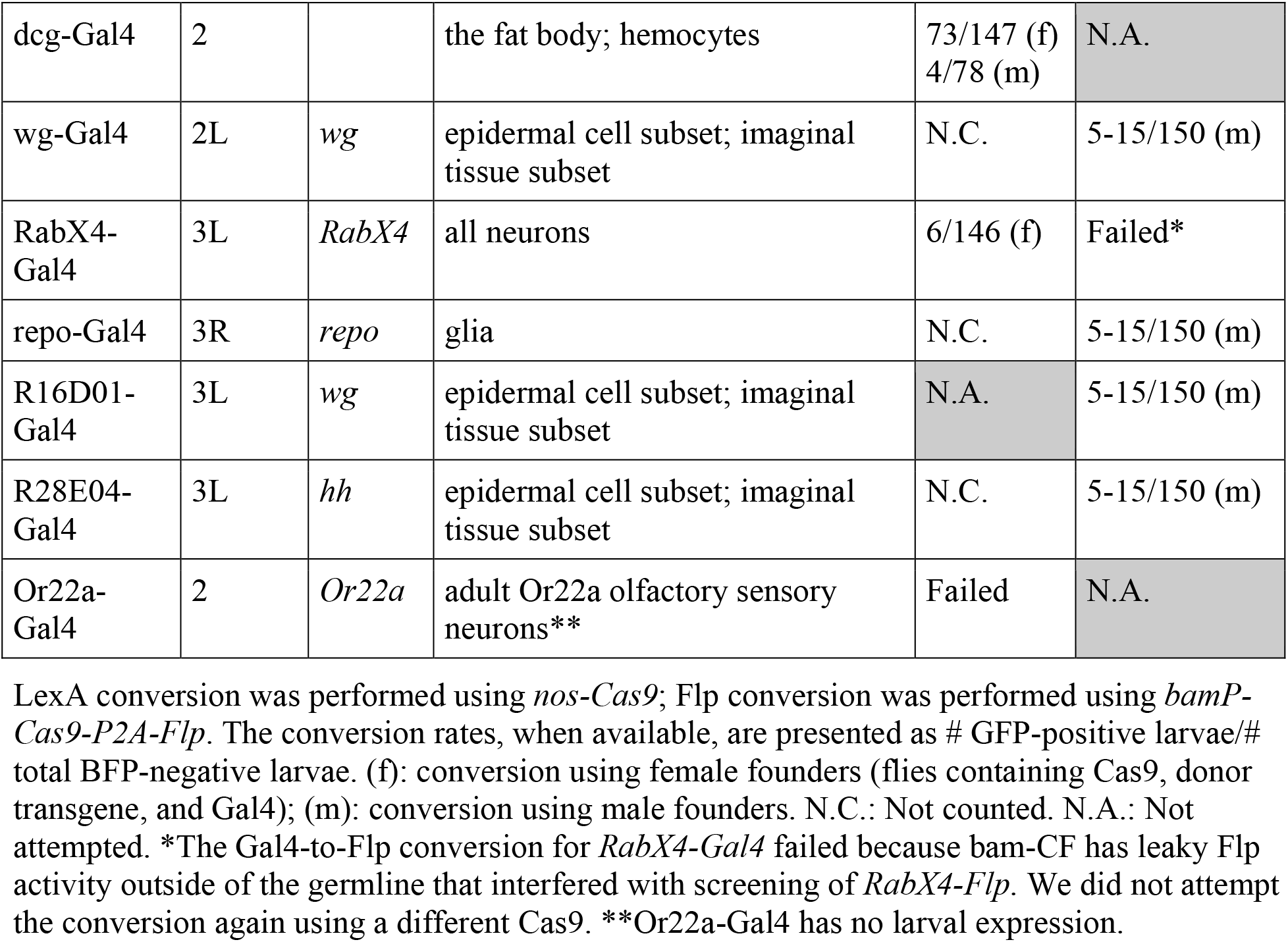
Summary of Gal4 lines and conversion rates.

We used donor transgenes located on the same chromosomes as the Gal4 insertions for conversion. For 14 out of 15 conversion experiments, we were able to identify larvae expressing the reporter in the expected pattern and to derive fly lines containing the converted chromosome from these larvae. Although the conversion frequency varied from experiment to experiment (Table 2), we recovered enough GFP-positive larvae by screening 100-300 candidates. The only exception was *RabX4-Gal4* to *RabX4-Flp* conversion, in which *bam-Cas9-P2A-Flp* (Chen *et al*.2020) produced leaky somatic Flp activity that interfered with the screening. In addition, we also tested Gal4-to-LexA conversion for *Or22a-Gal4* (Table 2), which has no larval expression but is expressed in a small number of olfactory neurons in the antenna. Because the adult cuticle is not transparent, *Or22a-Gal4* represents a challenging test case. We failed to detect obvious GFP signals in candidate adult flies using our setup.

### Comparison of the activity patterns of converted LexAGAD and Flp lines with those of the original Gal4 lines

To evaluate the activity patterns of the resultant LexAGAD and Flp lines, we crossed them to reporters (Table 1) and compared their activity patterns to those of their corresponding Gal4 lines. Cytosolic GFP reporters were used for lines that are expressed in non-neural tissues (Figure 3), while membrane-targeted GFP (mGFP) was used to examine the processes of neurons and glia (Figures 4A–4B’’). Nuclear GFP (nGFP) was used to locate the cell bodies of neurons (for OK371 and OK319) in the densely packed ventral nerve cord (VNC) (Figures 4C–4D’).

**Figure 3.**
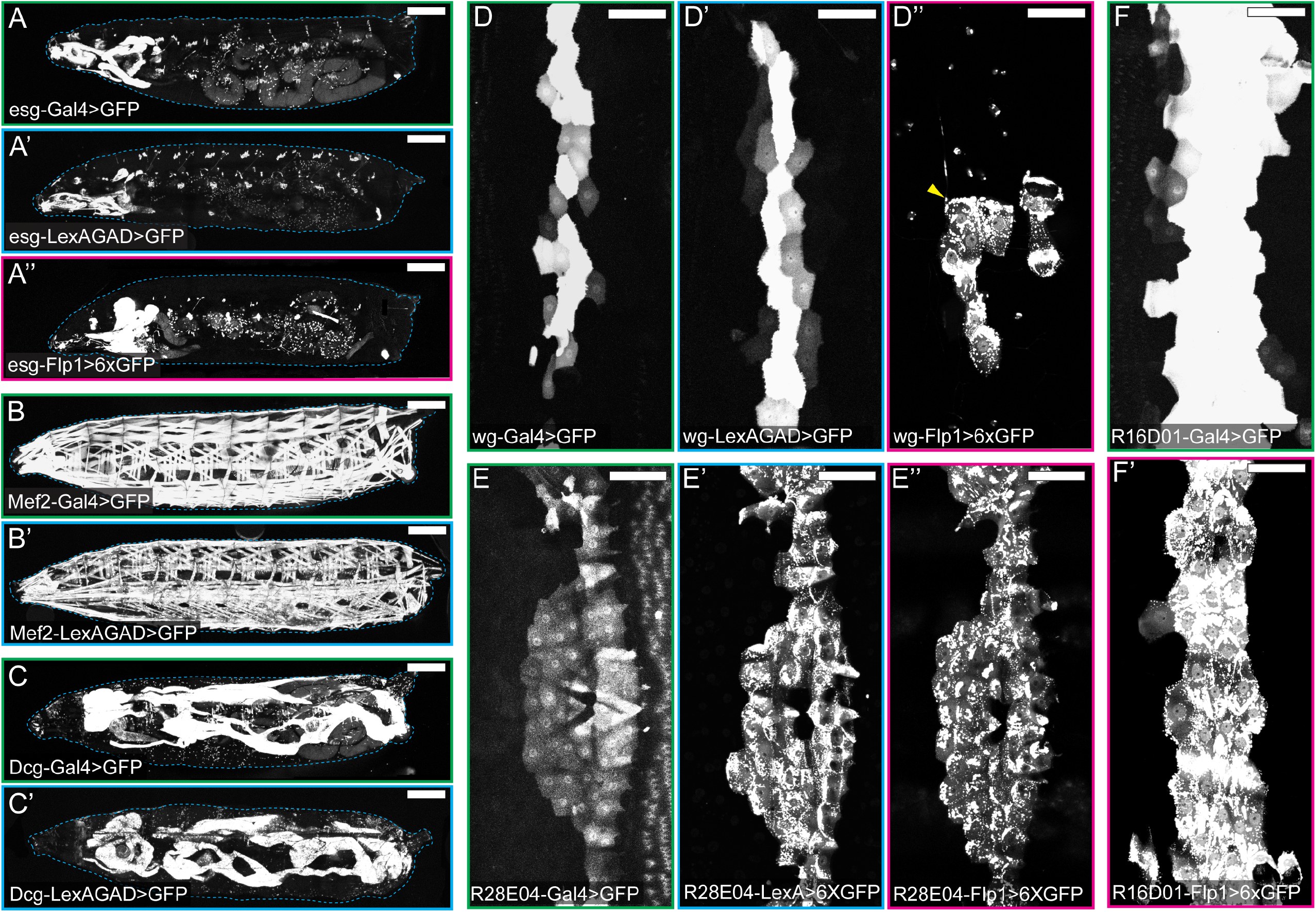
Comparison of Gal4 and converted LexA and Flp lines in non-neural tissues. (A-A”) Activity patterns of *esg-Ga14* (A), *esg-LexAGAD* (A’), and *esg-Flp1* (A”) in the whole larval body. (B-B’) Activity patterns of *Mef2-Ga14* (B) and *Mef2-LexAGAD* (B’) in the whole larval body. (C-C’) Activity patterns of *Dcg-Ga14* (C) and *Dcg-LexAGAD* (C’) in the whole larval body. (D-D”) Activity patterns of *wg-Gal4* (D), *wg-LexAGAD* (D’), and *wg-Flpl* (D”) in epidermal cells of a single hemisegment. Arrowhead (D”) indicates the cell body of a sensory neuron. (E-E”) Activity patterns of *R28E04-Ga14* (E), *R28E04-LexAGAD* (E’), and *R28E04-Flp1* (E”) in epidermal cells of a single hemisegment. (F-F”) Activity patterns of *R16D01-Gal4* (F) and *R16D01-LexAGAD* (F’) in epidermal cells of a single hemisegment. Refer to Table 1 for reporter lines used. Scale bar: 300 μm (A-C’); 100 μm (D-F’).

**Figure 4.**
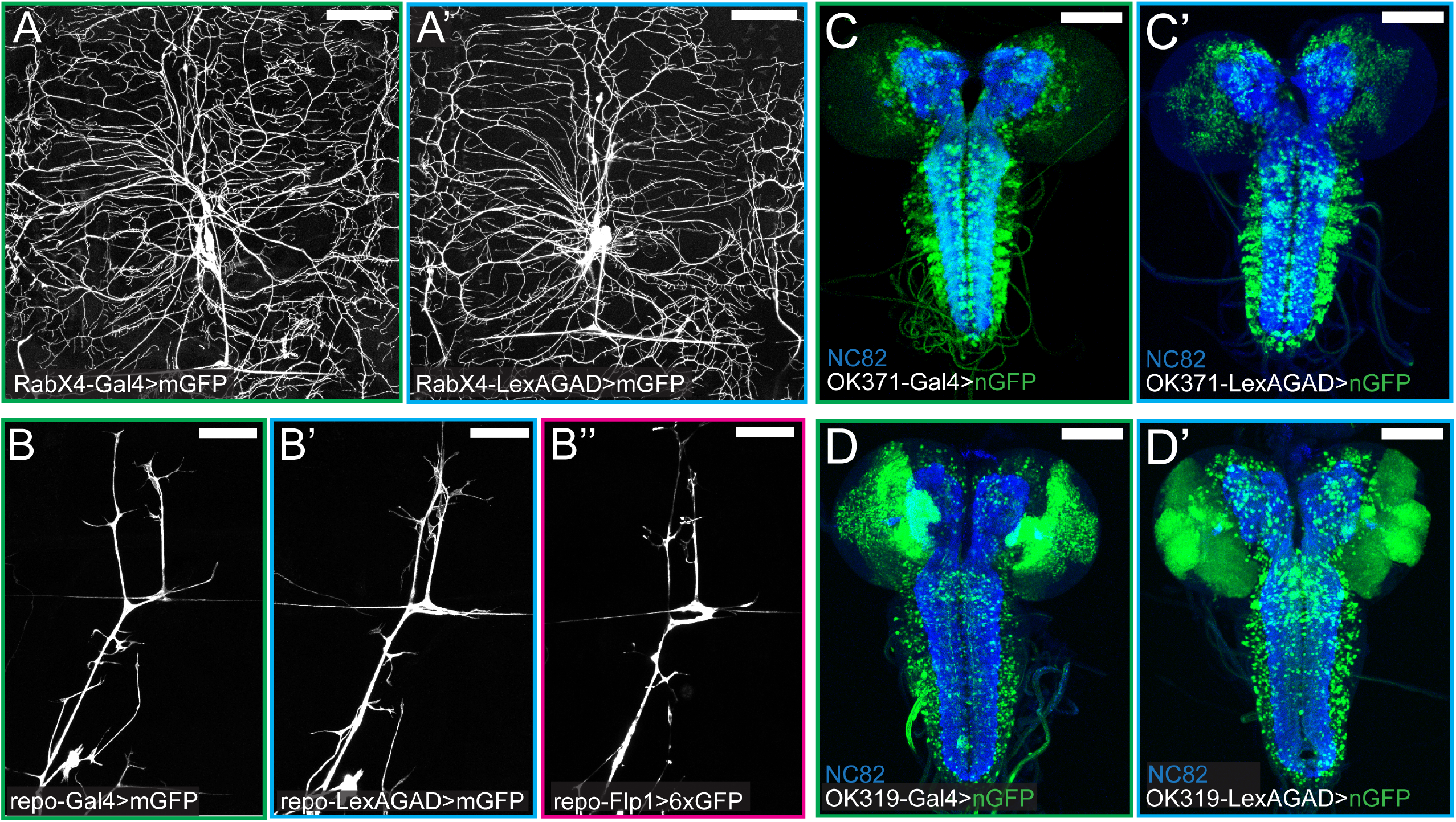
Comparison of Gal4 and converted LexA and Flp lines in the nervous system. (A-A’) Activity patterns of *RabX4-Gal4* (A) and *RabX4-LexAGAD* (A’) in a single dorsal hemisegment. (B-B’) Activity patterns of *repo-Gal4* (B), *repo-LexAGAD* (B’), and *repo-Flp1* (B”) in a single dorsal hemisegment. (C and C’) Activity patterns of *OK371-Gal4* (C) and *OK371-LexAGAD* (C’) in the larval brain. (D and D’) Activity patterns of *OK319-Gal4* (D) and *OK319-LexAGAD* (D’) in the larval brain. Refer to Table 1 for reporter lines used. Scale bar: 100 μm in all panels.

The activity patterns of the converted LexAGAD lines faithfully recapitulated the expression patterns of the corresponding Gal4 lines (Figures 3 and 4), with the only exception of *wg-LexAGAD*. Although *wg-LexAGAD* has a similar activity pattern as *wg-Gal4* in the larval epidermis (Figures 3D and 3D’), it showed broader activity in the wing pouch and restricted expression elsewhere in the wing imaginal disc of the late 3^rd^ instar larva as compared to *wg-Gal4* (Figures S2A and S2B). In comparison, two of the five converted Flp lines did not show identical activity patterns as their Gal4 counterparts. Specifically, *esg-Flp1* did not label all histoblasts but occasionally showed activity in some tracheal and muscle cells (Figure 3A”). Unlike *wg-Gal4* that is active in a narrow strip of epidermal cells along the dorsal-ventral axis of each hemisegment (Figure 3D), *wg-Flp1* labeled a smaller cluster of epidermal cells, as well as few peripheral neurons (arrowhead), in every hemisegment (Figure 3D”). In the wing imaginal disc, while *wg-Gal4* activity was detected at the dorsal/ventral boundary of the wing disc (Figure S1A), where *wg* expression is expected (Diaz-Benjumea and Cohen 1995;Kim *et al*. 1995), *wg-Flp1* resulted in labeling of distinct cell patches in dorsal and ventral compartments (Figure S2B). The discrepancies between Gal4 and Flp could be due to different thresholds required for activating their corresponding reporters and the fact that Flp patterns result from accumulation of activities throughout the developmental history while Gal4 patterns reflect current activity.

## Discussion

HACK is a convenient method for converting one genetic reagent to another through genetic crosses. With prebuilt donor transgenes, Gal4 can be converted into other reagents without needing cloning or injection, greatly simplifying the process required for generating new reagents. Converted reagents in theory should have similar activity patterns as the original Gal4 lines and thus, in most cases, need very little characterization. This method has been successfully used to convert Gal4 into QF, split Gal4, Gal80, and Cas9. In this study, we further expand the existing toolbox and make reagents available for generating tissue-specific LexAGAD and Flp lines from Gal4 lines. This conversion process is straightforward and can be performed in any *Drosophila* lab that is equipped with a fluorescence dissecting microscope. The ability to expand the current choices of LexA and Flp reagents to match those available for Gal4 will provide fly researchers greater flexibility for studying their questions.

Our HACK method differs from other similar approaches in the design of the donor constructs. The first important difference is that we used the CR7T-U63(2.1) design for expressing dual gRNAs. This design is specifically optimized for the *Drosophila* germline (Koreman *et al*. 2021). With higher mutagenic efficiency in the germline, this design is predicted to improve the conversion rate. Second, instead of using the 3xP3-RFP marker for selecting potential convertants (Lin and Potter 2016), we rely on LexA- or Flp-dependent reporter expression as the primary means for identifying the converted chromosomes. Although 3xP3-RFP is more convenient to screen in adults because of RFP expression in the eye, it has some disadvantages. Because 3xP3-RFP will be carried over into the converted reagents, it may interfere with subsequent experiments and, in most cases, needs to be first removed by the Cre recombinase. Also, incomplete homologous recombination events can result in false positive candidates that have incorporated 3xP3-RFP marker but not functional converted reagents (Chen *et al*. 2020). In comparison, screening based on reporter expression directly identifies correctly converted chromosomes and thus does not need additional validation by genomic PCR. The converted reagents can be directly used in subsequent experiments. The additional ubi-nlsBFP marker serves as a selection maker for distinguishing the donor chromosome but is not absolutely needed.

Screening KI events based on expression patterns also has some caveats. When Gal4 expressing cells are too sparse or buried too deeply inside the body, especially in adults that have opaque cuticles, the fluorescence from the expressing cells may not be distinguishable for screening. For example, we failed to convert Or22a-Gal4, which is only expressed in a small number of olfactory neurons whose cell bodies are buried inside the antenna. In situations like this, donor templates that incorporate visible selection markers may still be better choices for the conversion.

As reported previously (Lin and Potter 2016), the frequency of conversion can vary greatly among different Gal4 lines, likely due to the local chromatin conformation. We noticed a wide range of conversion efficiencies as well (Table 2). While *OK371-Gal4* and *dcg-Gal4* were very easy to convert, *RabX4-Gal4* and *FlyLight* Gal4 lines were more refractory to conversion. For Gal4 lines that are difficult to convert, the efficiency can be improved by taking several measures. First, when *nos-Cas9*(Port *et al*. 2014) is used as the germline Cas9, we found that female founders, which contain Cas9, Gal4, and the donor transgene, gave higher conversion rates than male founders. Second, *bam-Cas9-P2A-Flp*, which is expressed in germline precursor cells but not in germline stem cells, was reported to perform better in germline HDR (Chen *et al*.2020). Our preliminary comparisons support this conclusion. Thirdly, even though we have not confirmed it, using two copies of donor transgenes may improve efficiency as well. The Lee group recently reported the E-Golic+ for genetic cross-based KI (Chen *et al*. 2020). This system incorporates *bam-Cas9-P2A-Flp* and uses induced lethality to eliminate non-converted chromosomes and thus could dramatically improve the efficiency. Although this system also requires removing selection markers from positive candidates, it may still be a better choice for converting Gal4 insertions extremely difficult to convert by other means.

Besides HACK, new LexA and Flp reagents can also be generated by other means. For example, MiMIC (Venken *et al*. 2011) and CRIMIC (Lee *et al*. 2018) lines, for which large collections are available, can be converted into different effectors using appropriate Recombinase Mediated Cassette Exchange (RMCE) donor lines. The InSITE system (Gohl *et al*.2011) also allows for effector conversion of >1,300 enhancer-trap Gal4 lines based on RMCE. Although donor lines for converting these resources into LexA (except for InSITE) and Flp reagents still need to be established, these systems offer complementary approaches for expanding LexA and Flp choices. Lastly, although the enhancer-fusion Gal4 lines in the FlyLight (Jenett *et al*. 2012) and VT (Kvon *et al*. 2014) collections are compatible with HACK, we found that these Gal4 transgenes inserted into the attP2 site are relatively more difficult to convert by HACK. Because the enhancer sequence for each of these Gal4 lines is molecularly defined, making and transforming new enhancer-fusion constructs may be a more reliable approach for generating corresponding LexA and Flp strains (Poe *et al*. 2017).

The HACK method can in principle be used to convert Gal4 into any other type of genetic reagent. Although we present tools for generating LexA and Flp in this study, our donor vectors can be modified for conversion of many other types of reagents, such as GeneSwitch-Gal4 (Osterwalder *et al*. 2001), LexA::P65 (Pfeiffer *et al*. 2010), cpf1 (Zetsche *et al*. 2015), etc.

LexA conversion was performed using *nos-Cas9*; Flp conversion was performed using *bamP-Cas9-P2A-Flp*. The conversion rates, when available, are presented as # GFP-positive larvae/# total BFP-negative larvae. (f): conversion using female founders (flies containing Cas9, donor transgene, and Gal4); (m): conversion using male founders. N.C.: Not counted. N.A.: Not attempted. *The Gal4-to-Flp conversion for *RabX4-Gal4* failed because bam-CF has leaky Flp activity outside of the germline that interfered with screening of *RabX4-Flp*. We did not attempt the conversion again using a different Cas9. **Or22a-Gal4 has no larval expression.

## Data Availability

The donor vectors are available at Addgene: pHACK(Gal4)-DONR(T2A-LexAGAD) (Addgene # 194769); pHACK(Gal4)-DONR(T2A-Flp1) (Addgene # 194770). Other plasmids are available upon request. *Drosophila* strains are available at Bloomington *Drosophila* Stock Center or upon request. The authors affirm that all data necessary for confirming the conclusions of the article are present within the article, figures, and tables.

## Acknowledgments

We thank Tzumin Lee, Bloomington *Drosophila* Stock Center (NIH P40OD018537), and KYOTO Stock Center for fly stocks; Dion Dickman and members of Han lab for critical reading and suggestions on the manuscript. This work was supported by NIH grants (R01NS099125, R21OD023824, and R24OD031953) awarded to C.H..

## Author Contributions

S.K., A.T.Y., B.W., and C.H. designed research; S.K., A.T.Y., B.W., and P.F.G. performed research; B.W. contributed new reagents/analytic tools; S.K., A.T.Y., and C.H. analyzed data; C.H. wrote the manuscript; S.K., A.T.Y., and C.H. edited the manuscript; C.H. acquired funding.

## Figure Legend

**Figure S1. Validation of converted LexAGAD and Flp lines by genomic PCR**

(A) Diagram of converted Flp transgene and genomic PCR results for *esg*, *repo*, and *wg* lines. The positions of 5’ and 3’ homology arms (HAs), binding locations of PCR primers, and expected sizes of PCR products are indicated in the diagram. The DNA gel shows PCR results of the Flp donor line, the original Gal4 (1), and the converted Flp (2).

(B) Diagram of converted LexAGAD transgene and genomic PCR results for *repo, OK371*, and *wg* lines. The diagram and PCR results are labeled similarly to (A).

The primers used for PCR amplifications are (a) CTTGAAGCAAGCCTCCTGAAAG; (b) AGTGGTATTAAACATCCCTGTAGTG; (c) TGACGCACCAACACCTTTG; (d) CAGGAGGTTCTGGATTACCTGAG; (e) GAGAGCCTTCATTGGATCTTCTAC; (f) ACCATCTACCACGGTATCATTGAG.

**Figure S2. Comparison of *wg-Gal4, wg-LexAGAD*, and *wg-Flp* lines in the wing disc**

(A-C) Activity patterns of *wg-Gal4* (A), *wg-LexAGAD* (B), and *wg-Flp1* (C) in a 3^rd^ instar wing imaginal disc. GFP is in green; DAPI staining is in blue.

Refer to Table 1 for reporter lines used. Scale bar: 100 μm in all panels.

